# Hsp70 chaperons TDP-43 in dynamic, liquid-like phase and prevents it from amyloid aggregation

**DOI:** 10.1101/2020.12.29.424685

**Authors:** Jinge Gu, Chen Wang, Rirong Hu, Yichen Li, Shengnan Zhang, Yunpeng Sun, Qiangqiang Wang, Dan Li, Yanshan Fang, Cong Liu

**Author notes:** These authors contributed equally (Jinge Gu, Chen Wang, Rirong Hu). **Correspondence to:** Yanshan Fang, Cong Liu.

## Abstract

TAR DNA binding protein 43 kDa (TDP-43) undergoes liquid-liquid phase separation (LLPS) and forms reversible, cytoprotective nuclear bodies (NBs) in response to stress in cells. Abnormal liquid-to-solid phase transition condenses TDP-43 into irreversible pathological fibrils, which is associated with neurodegenerative disorders including amyotrophic lateral sclerosis (ALS) and frontotemporal degeneration (FTD). However, the mechanisms how cells maintain the dynamics of TDP-43 NBs in stressed conditions are not well understood. Here, we show that the molecular chaperon heat shock protein 70 (Hsp70) is recruited into TDP-43 NBs in stressed cells. It co-phase separates with TDP-43 and delays the maturation of TDP-43 droplets *in vitro*. In cells, downregulation of Hsp70 not only diminishes the formation but also reduces the dynamics of TDP-43 NBs especially during prolonged stress, which potentiates the cytotoxicity of TDP-43. Using NMR, we reveal that Hsp70 binds to the highly aggregation-prone, transient α-helix of TDP-43 via its nucleotide-binding domain, which keeps TDP-43 in the highly dynamic, liquid-like phase and prevents pathological aggregation of TDP-43 both *in vitro* and in cells. Collectively, our findings demonstrate a crucial role of Hsp70 in chaperoning TDP-43 in the liquid-like phase, which provides a novel layer of the molecular mechanism how chaperons help proteins to remain functional and protect cells from stressed and/or diseased conditions.

## INTRODUCTION

TDP-43 is a nuclear RNA-binding protein (RBP) that plays a pivotal role in RNA processing and homeostasis (Birsa et al., 2020; Lee et al., 2011; Zhao et al., 2018). In response to stress, TDP-43 undergoes dynamic and reversible LLPS, which is involved in the formation of different membraneless organelles in the cell such as stress granules (SGs) and NBs (Li et al., 2013; McGurk et al., 2018; Wang et al., 2020). Chronic stress and aging may cause aberrant phase transition and the formation of irreversible amyloid aggregates of TDP-43 in the nucleus and cytoplasm, which is associated with a variety of neurodegenerative diseases including ALS, FTD and Alzheimer’s disease (AD) (Chen-Plotkin et al., 2010; Ramaswami et al., 2013; Shukla and Parker, 2016; Fahrenkrog and Harel, 2018; Gasset-Rosa et al., 2019).

In a recent study, we revealed that TDP-43 forms dynamic, liquid-like NBs that mitigate the cytotoxicity in response to various conditions of cellular stresses. This process is mediated by the long non-coding RNA (lncRNA) *Nuclear Enriched Abundant Transcript 1* (*NEAT1*), which promotes TDP-43 phase separation and is required for the assembly of TDP-43 NBs in stressed cells (Wang et al., 2020). Other types of RNAs such as tRNA suppress the LLPS of TDP-43 and other disease-related RBPs (Maharana et al., 2018; Mann et al., 2019). Stress upregulates *NEAT1* to trigger the LLPs of TDP-43 NBs in the general suppressive environment of the nucleoplasm. In the meanwhile, TDP-43 is highly aggregation-prone protein. Thus, it is important and challenging for cells not to over-condense TDP-43 NBs and to keep them dynamic and reversible in order to survive stress and regain function once stress is released. Nevertheless, the mechanism by which cells maintain the highly aggregation-prone TDP-43 in the liquid-like phase and prevent TDP-43 NBs from aggregation under stressed conditions remains elusive.

Molecular chaperones play a crucial role in maintaining protein homeostasis in cells (Hartl et al., 2011). For example, heat shock proteins (Hsp) such as Hsp70, Hsp40 and Hsp60 safeguard proteins against stress-induced misfolding and aggregation (Hartl, 1996). In particular, Hsp70 is closely associated with the regulation of TDP-43 proteostasis – (1) Hsp70 was found to co-exist with TDP-43 in SGs (Jain et al., 2016); (2) Hsp70 can interact with TDP-43 in mammalian cell cultures (Freibaum et al., 2010; Udan-Johns et al., 2014; Kitamura et al., 2018); (3) increase of Hsp70 can suppress TDP-43-mediated toxicity in fly models (Estes et al., 2011; Coyne et al., 2017); and more importantly (4) Hsp70 expression in the spinal cord of sporadic ALS patients with TDP-43 aggregates is significantly decreased (Chen et al., 2016), suggesting that dysregulation of Hsp70 may be involved in pathological aggregation of TDP-43 in ALS and related diseases.

In this study, we show that Hsp70 protein translocates to the nucleus and is co-localized with TDP-43 NBs in response to stress. Further investigation reveals that Hsp70 co-phase separates with TDP-43 *in vitro* and it promotes the assembly of TDP-43 NBs in cells. The presence of Hsp70 in TDP-43 NBs helps to maintain them in the dynamic, liquid-like phase, which increase the cell viability in stressed conditions. Hsp70 executes this function mainly via is N-terminal domain (NTD) that binds to the hydrophobic and aggregation-prone region of TDP-43, which stabilizes TDP-43 in the highly dynamic, liquid-like phase and prevents it from abnormal liquid-to-solid transition and pathological aggregation. Together, the finding of the participation and function of Hsp70 in TDP-43 NBs provides a new paradigm how cells maintain the aggregation-prone RBPs such as TDP-43 in condensed but highly dynamic NBs in response to stress.

## RESULTS

### Hsp70 co-localizes with stress-induced TDP-43 NBs in cells and can co-phase separate with TDP-43 *in vitro*

In a recent study, we reported that TDP-43 formed cytoprotective NBs in various conditions of cellular stress (Wang et al., 2020). In this work, we set out to examine whether molecular chaperons play a role in the regulation of TDP-43 NBs. Since Hsp70 was previously shown to co-localize with TDP-43 in SGs (Jain et al., 2016), we asked whether Hsp70 was also associated with TDP-43 in NBs. As shown in Fig. 1a-b, red fluorescent protein (RFP)-Hsp70 expressed in HeLa cells was mostly diffused and predominantly localized in the cytoplasm, which was consistent with the expression pattern of endogenous Hsp70 protein as examined by immunocytochemistry (Fig. S1a). Upon arsenic stress, Hsp70 displayed increased fluorescent signal in nucleus and was co-localized with TDP-43 NBs (Fig. 1c-d). Of note, the recruitment of Hsp70 into the TDP-43 NBs could be visualized by monitoring the fluorescence of the RFP tag but not by immunostaining with the anti-Hsp70 antibody (Fig. S1b), likely because the antibody could not penetrate into NBs as also reported by Yu, H., et al. (2020). This unfortunately excluded the possibility to unambiguously demonstrate the co-localization of endogenous Hsp70 with TDP-43 NBs. Nevertheless, as an additional layer of control, we expressed RFP in HeLa cells, which was diffused and located in both the nucleus and the cytoplasm (Fig. 1e-f), but it did not co-localize with TDP-43 NBs upon the arsenite treatment (Fig. 1g-h). This result confirmed that the recruitment of RFP-Hsp70 into TDP-43 NBs in response to stress was specific and not merely because Hsp70 was overexpressed.

**Fig. 1:**
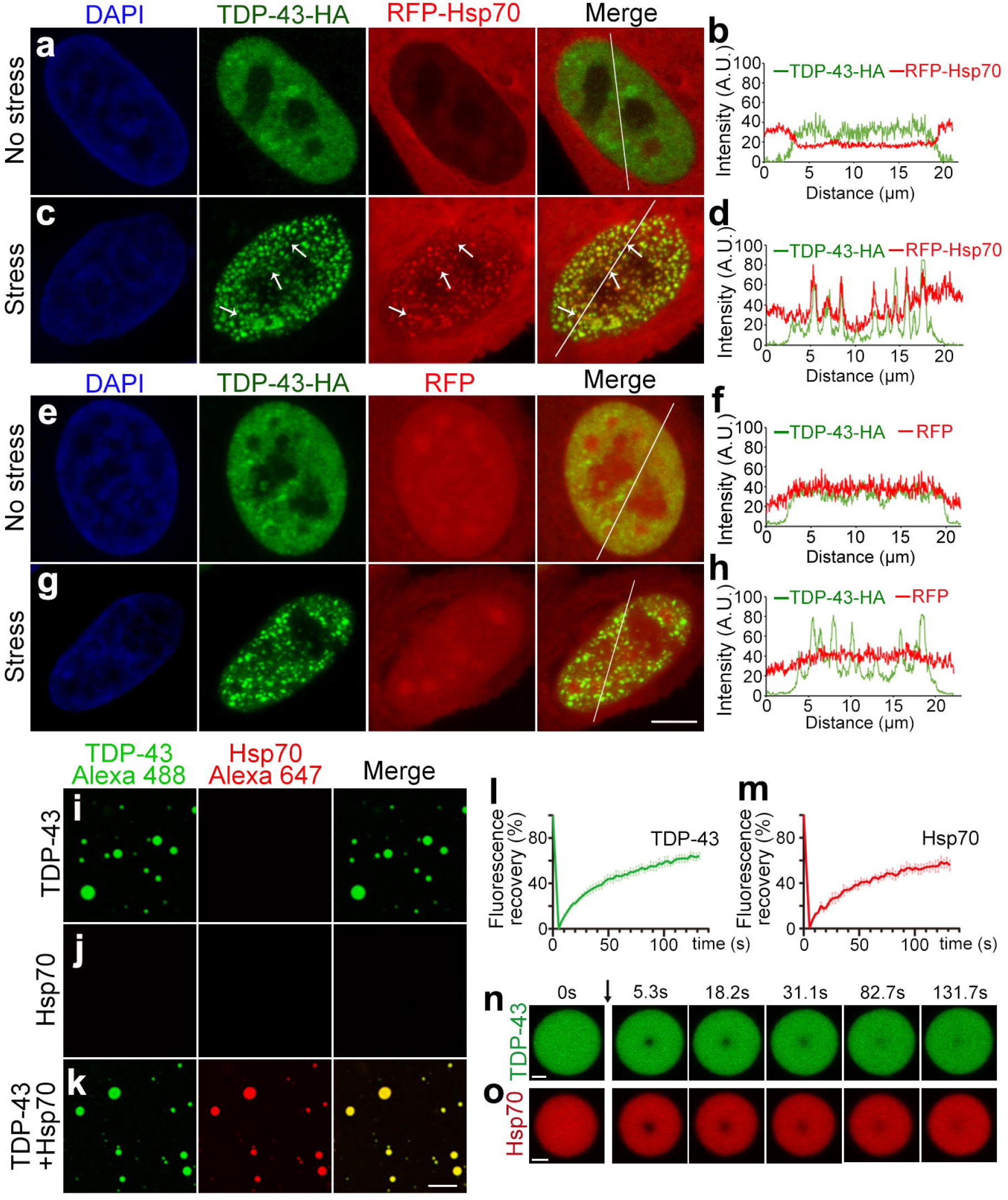
Hsp70 co-localizes with TDP-43 NBs in stressed cells and co-phase separates with TDP-43 *in vitro*. **a-h** Representative images (a, c, e, g) and the intensity profiles along the indicated line (b, d, f, h) of HeLa cells expressing RFP-Hsp70 (a-d) or RFP (e-h) together with TDP-43-HA in the absence or presence of cellular stress (250 μM of arsenite, 30 min). DAPI, nuclear labeling; anti-HA for TDP-43-HA; arrows, co-localization of Hsp70 with TDP-43 NBs. **i-k** Representative confocal images showing the *in vitro* LLPS of TDP-43-MBP (Alexa Fluor 488, green) alone (i), Hsp70 (Alexa Fluor 647, red) alone (j), and them together (k). The concentration of each component in the *in vitro* LLPS assay: 50 mM Tris-HCl, pH 7.5, 150 mM NaCl, 15% Dextran 70, 50 μM TDP-43-MBP and 10 μM Hsp70. **l-o** The FRAP analyses (l-m) and images of representative droplets (n-o) of TDP-43 (l, n, green) and Hsp70 (m, o, red) in the co-phase separated droplets in k. The black arrow indicates the photobleaching. Data shown are mean ± SEM, n = 5 (l and m). Scale bars, 5 μm in (a, c, e and g), 10 μm in (i-k) and 1 μm in (n-o).

As Hsp70 was co-localized with TDP-43 in stress-induced NBs in cells, we then examined whether Hsp70 could co-phase separate with TDP-43 in the *in vitro* LLPS assay. Full-length (FL) TDP-43 and Hsp70 proteins were purified and fluorescently labeled with Alexa 488 and Alexa 647, respectively. As previous reported (McGurk et al., 2018; Wang et al., 2018; Wang et al., 2020), TDP-43 underwent LLPS and formed liquid droplets in the *in vitro* de-mixing system (Fig. 1i). Hsp70 did not phase separate on its own at this condition (Fig. 1j). However, Hsp70 co-phase separated with TDP-43 in the droplets when mixed together with TDP-43 (Fig. 1k). Further, the fluorescence recovery after photobleaching (FRAP) assay showed that the intensity of the TDP-43 fluorescence signal rapidly recovered to over 60% within 130 s after photobleaching, and the FRAP of Hsp70 also exhibited similar recovery curve (Fig. 1l-o). This indicated that the co-phase separated droplets of TDP-43 and Hsp70 were highly dynamic and liquid-like. Together, these data indicated that Hsp70 could co-phase separate with TDP-43 *in vitro* and was recruited into TDP-43 NBs in stressed cells.

### Knockdown (KD) of Hsp70 dysregulates TDP-43 NBs and potentiates the cytotoxicity in prolonged stress

To understand the physiological significance of co-localization and co-phase separation of Hsp70 with TDP-43 NBs, we sought to examine the impact of downregulation of Hsp70 on TDP-43 NBs. Hsp70 protein is abundant in cells and the mammalian Hsp70 family has multiple members (Tavaria et al., 1996; Bettencourt and Feder, 2002; Nikolaidis and Nei, 2004; Brocchieri et al., 2008). We thus examined the basal mRNA expression levels of the main Hsp70 genes in HeLa cells (Fig. S2a) and their changes in response to arsenic stress (Fig. S2b). Among them, *HSPA8* (which encodes Hsc70) appeared to the most abundantly expressed, accounting for ~62% of total Hsp70 mRNAs in normal conditions (Fig. S2a); whereas in response to stress, *HSPA1A* (which encodes Hsp70) increased the most dramatically, displaying an upregulation of over 18 folds (Fig. S2b). Of note, we confirmed that the most abundant Hsp70 member, Hsc70, could also co-phase separate with TDP-43 *in vitro* and was co-localized with stress-induced TDP-43 NBs in cells (Fig. S3).

Next, we knocked down Hsp70 and Hsc70 using small interference RNA (siRNA) of *HSPA1A* and *HSPA8* (Fig. 2a-d; for simplicity, shown as si-Hsp70s). Strikingly, both the percentage of cells forming TDP-43 NBs (Fig. 2e) and the average counts of TDP-43 NBs per cell (Fig. 2f) in response to stress were significantly reduced in cells treated with si-Hsp70s compared to the control cells treated with scrambled siRNA (si-Ctrl). In contrast, si-Hsp70s did not significantly affect the formation of SGs as shown by the SG marker G3BP (Fig. 2g). And, the western blotting assay confirmed that KD of *HSPA1A* and *HSPA8* by siRNAs could reduce the total Hsp70 protein levels by about half (Fig. 2h-i).

**Fig. 2:**
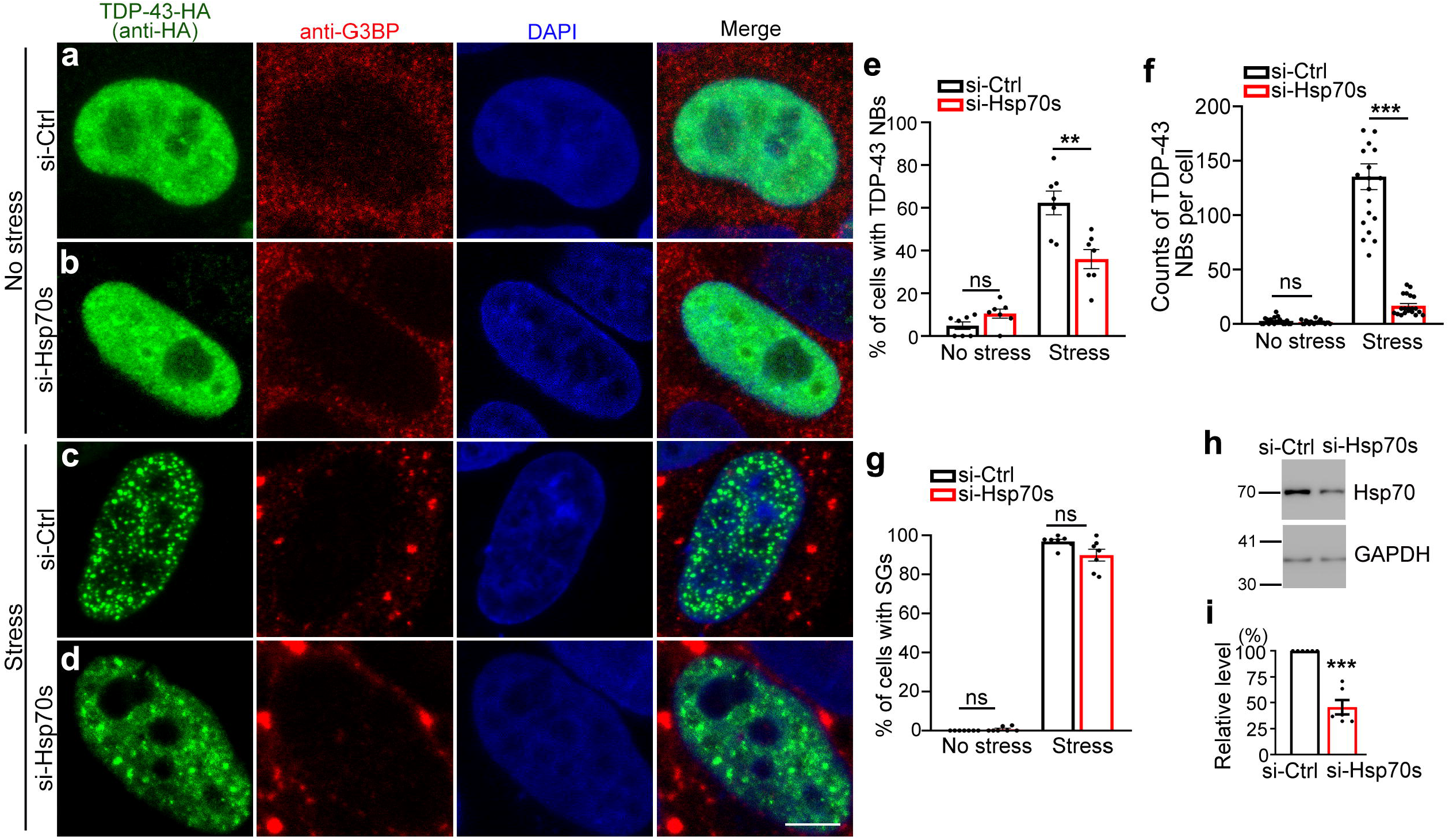
KD of Hsp70 reduces the assembly of TDP-43 NBs. **a-d** The effect of si-Hsp70s (both siRNAs against *HSPA1A* and *HSPA8*) on the assembly of stress-induced TDP-43 NBs compared to si-Ctrl (scrambled siRNA). Representative images of HeLa cells expressing TDP-43-HA in the absence (a-b) and presence (c-d) of arsenite (250 μM, 30 min) are shown. DAPI, nucleus; anti-G3BP, SGs. **e-g** The percentage of cells forming TDP-43 NBs (e), the average count of TDP-43 NBs per cell (f), and the percentage of cells forming SGs (g) in a-d are quantified. **h-i** The KD efficiency of si-Hsp70s is examined by western blotting analysis (h) and quantified in (i). GAPDH is used as a loading control. Mean ± SEM; n = ~80 cells per group in (e-f) and n = 20 cells in (g) from pooled results of 3 independent repeats, and n = 6 in (i). Statistical significance was determined by Student’s *t* test at *\*\*p* < 0.01 and \*\*\**p* < 0.001; ns, not significant. Scale bar, 5 μm.

We then examined how si-Hsp70s affected the dynamics of TDP-43 NBs by the FRAP assay (Fig. 3a-j). Interestingly, KD of Hsp70 did not alter the liquid-like, dynamic feature of TDP-43 NBs with a transient stress of 30-min arsenite treatment (Fig. 3b-d). However, when the arsenic stress persisted to 60 min, TDP-43 NBs in the cells with si-Hsp70s showed marked reduction of dynamics (Fig. 3e-g). And this was further worsened with prolonged stress of 120-min arsenite treatment, as the enhanced green fluorescence protein (EGFP) signal of EGFP-TDP-43 in the NBs recovered to ~50% within 250 s after photobleaching in the control cells while that of EGFP-TDP-43 NBs in the si-Hsp70s cells hardly recovered (Fig. 3h-j). As a result, we found that KD of Hsp70 made the cells more vulnerable to TDP-43-induced cytotoxicity and showed significantly decreased cell viability under prolonged cellular stress (Fig. 3k). Overexpression (OE) of Hsp70 did not show a remarkable effect on TDP-43 NBs or cell viability under stress (Fig. S4). Thus, although not sufficient, the recruitment of Hsp70 to TDP-43 NBs was required for the assembly and sustaining of the highly dynamic, liquid-like property of TDP-43 NBs during cellular stress.

**Fig. 3:**
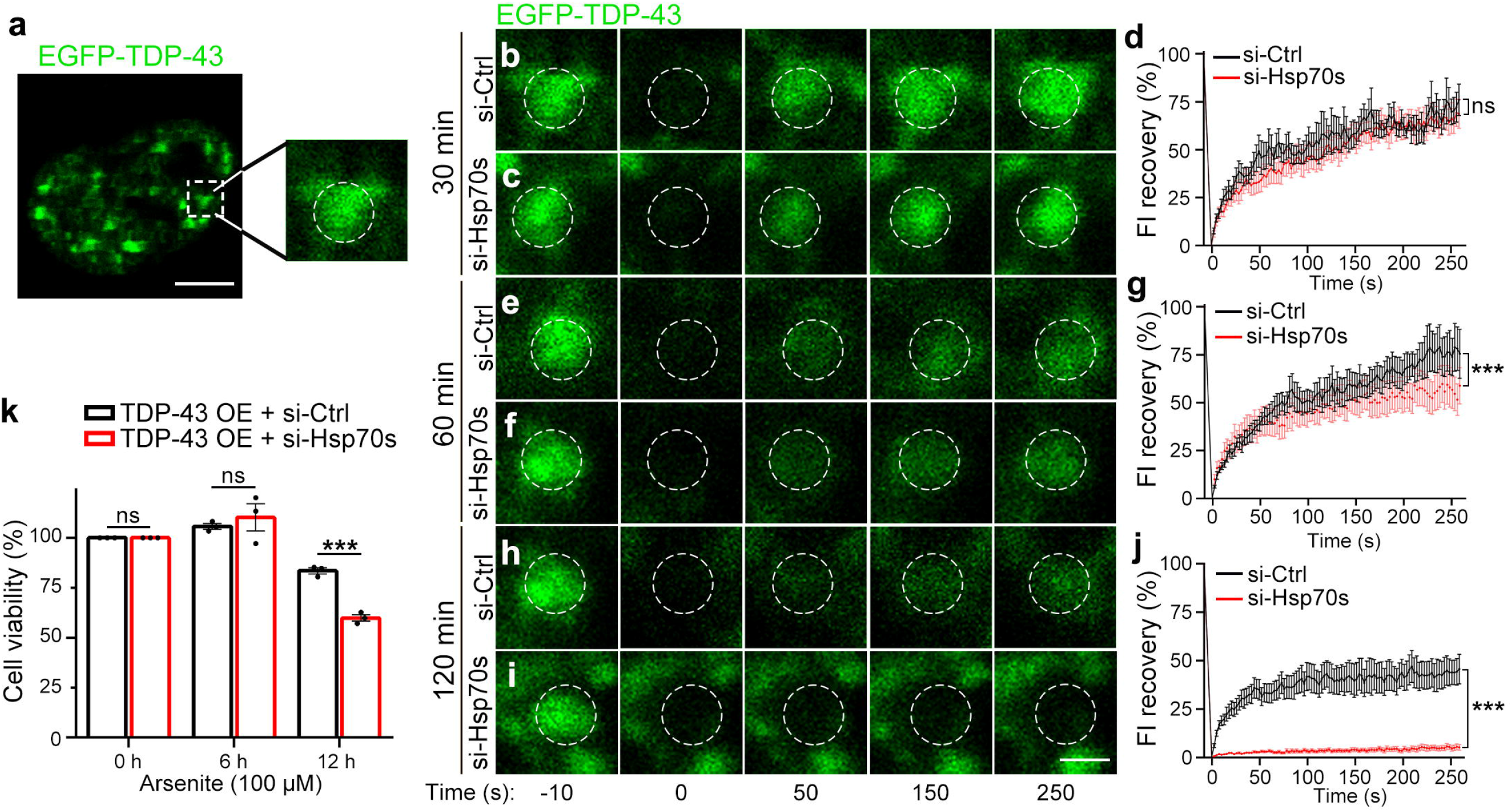
KD of Hsp70 accelerates liquid-to-solid maturation of TDP-43 NBs and potentiates the cytotoxicity in prolonged cellular stress. **a** A representative image showing stress-induced EGFP-TDP-43 NBs in living HeLa cells and the region (dashed line) subject to photobleaching in the FRAP assay in b-j. **b-j** Representative images of EGFP-TDP-43 NBs (b, c, e, f, h, i) and the fluorescent intensity (FI) recovery curves (d, g, j) of the FRAP assays. Cells transfected with scrambled siRNA (si-Ctrl) or siRNAs against *HSPA1A* and *HSPA8* (si-Hsp70s) are treated with arsenite (250 μM) for 30 min (b-d), 60 min (e-g) or 120 min (h-j) as indicated. **k** The viability of HeLa cells transfected with TDP-43-HA together with si-Ctrl or si-Hsp70 under stress (100 μM) for different durations as indicated is assessed using the CCK-8 assay. Mean ± SEM; n = 8 in (d, g, j) and n = 3 in (k). Two-way ANOVA (d, g, j), Student’s *t* test (k); \*\*\**p* < 0.001; ns, not significant. Scale bars, 5 μm in (a) and 1 μm in (b-i).

### Hsp70 chaperones TDP-43 in the dynamic, liquid-like phase

We went further to investigate the role of the co-LLPS of Hsp70 with TDP-43 and the underlying mechanism by which the recruitment of Hsp70 helped TDP-43 NBs to maintain in the liquid-like phase. Purified TDP-43 protein formed liquid-like droplets *in vitro*, which matured along the time as the FRAP assays indicated that the dynamics of the TDP-43 droplets dropped dramatically after 40 min of *in vitro* incubation (Fig. 4a). Strikingly, addition of purified Hsp70 protein in the *in vitro* LLPS system significantly delayed the maturation process, as the TDP-43 droplets with the presence of Hsp70 showed much faster and higher fluorescence recovery after photobleaching (Fig. 4a).

**Fig. 4:**
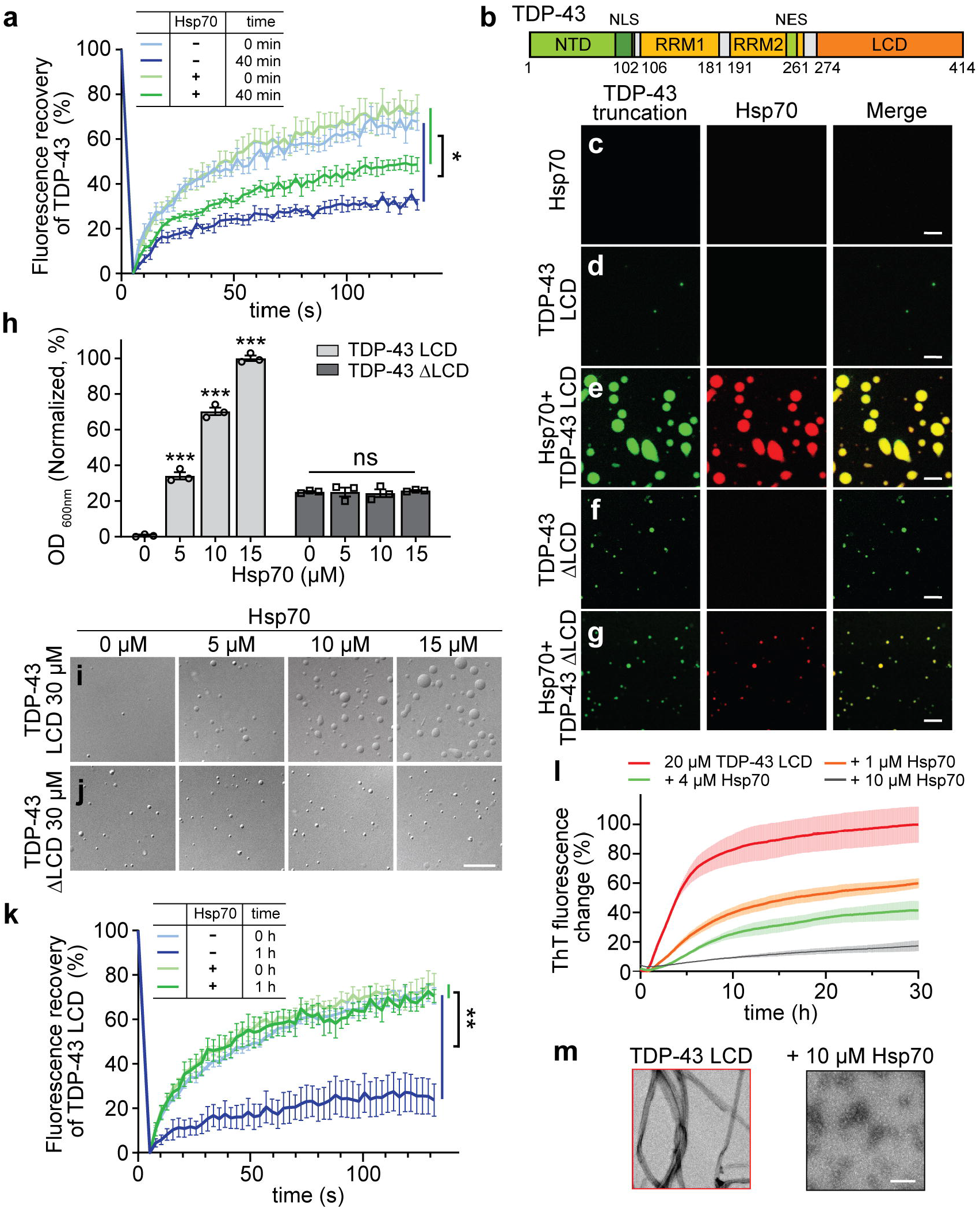
Hsp70 maintains TDP-43 in dynamic liquid-droplets and prevents it from amyloid aggregation. **a** FRAP analyses of TDP-43 liquid-droplets formed in the *in vitro* LLPS assay (50 μM TDP-43-MBP, 150 mM NaCl, pH 7.5, 15% Dextran 70) in the absence or presence of Hsp70 (75 μM). The FRAP assay was performed right after mixing the proteins together (0 min) or after co-incubation for 40 min. **b** Schematic of the TDP-43 domains. NTD, N-terminal domain; RRM, RNA recognition motif; LCD, low-complexity domain; NLS, nuclear localization signal; NES, nuclear export sequence. **c-g** Co-LLPS of Hsp70 (red) with TDP-43 LCD or TDP-43 ΔLCD (green). Fluorescence images show Hsp70 (15 μM) alone (c), TDP-43 LCD (30 μM) alone (d) or mixed with Hsp70 (15 μM) (e) in 50 mM NaCl, 20 mM MES, pH 6.0, and TDP-43 ΔLCD (30 μM) alone (f) or mixed with Hsp70 (15 μM) (g) in 50 mM NaCl, 20 mM Tris-HCl, pH 7.5. **h-j** The turbidity measurement (h) and representative DIC images (i-j) of TDP-43 LCD or ΔLCD (30 μM) with increasing concentrations of Hsp70 as indicated. The other conditions are same as in (c-g). **k** The FRAP assay of the LLPS liquid droplets of TDP-43 LCD (60 μM) in the absence or presence of Hsp70 (60 μM) at 0 h or 1 h after mixing (100 mM NaCl, pH 7.5). **l** The ThT fluorescence assay of TDP-43 LCD (20 μM) with different concentrations of Hsp70 as indicated. **m** Negative-staining TEM images of the ThT samples at 30 h in (l). Mean ± SEM; n = 6 in (a), n = 3 in (h), n = 5 in (k) and n = 3 in (l). Two-way ANOVA in (a, k), one-way ANOVA in (h); *\*p* < 0.05, \*\**p* < 0.01 and \*\*\**p* < 0.001, ns, not significant. Scale bars, 5 μm in (c-g), 10 μm in (i-j) and 200 nm in (m).

TDP-43 consists of an NTD, two RNA recognized motif (RRM), and a prion-like, low complexity domain (LCD) (Fig. 4b). To determine which region in the TDP-43 protein mediated the regulatory effect of Hsp70, we purified the truncated TDP-43 LCD and TDP-43 ΔLCD proteins. We found that Hsp70 could co-phase separate with both the LCD and ΔLCD of TDP-43 (Fig. 4c-g). However, Hsp70 significantly promoted the LLPS of TDP-43 LCD but only showed minimal effect on that of TDP-43 ΔLCD (Fig. 4h-j), suggesting that the LCD of TDP-43 played a significant role in mediating the interaction and co-LLPS of TDP-43 with Hsp70.

TDP-43 LCD was previously identified as the key region in mediating liquid to solid phase transition and pathological fibril formation of TDP-43 (Johnson et al., 2009; Babinchak et al., 2019; Zhuo et al., 2020). Indeed, TDP-43 LCD formed liquid-like droplets on its own, which underwent rapid maturation with dramatically decreased dynamics, recapitulating the maturation process of the droplets formed by FL TDP-43 (Fig. 4k). Strikingly, the FRAP assay showed that with Hsp70, the dynamics of the droplets of TDP-43 LCD were well maintained even after an hour of incubation (Fig. 4k). Thus, Hsp70 was a potent “dynamics keeper”, whose presence in the droplets of TDP-43 LCD effectively prevented them from maturation. To further examine whether Hsp70 could prevent pathological amyloid fibrillation of TDP-43 LCD, we conducted Thioflavin T (ThT) fluorescence kinetic assay combined with negative-staining transmission electron microscopy (TEM). As shown in Fig. 4l-m, TDP-43 LCD spontaneously formed amyloid fibrils, which were inhibited by Hsp70 in a dose-dependent manner. Together, our results demonstrated that Hsp70 not only co-phase separated with TDP-43, but also stabilized it in a liquid-like state by preferentially interacting with the LCD of TDP-43, which prevented the amyloid fibrillation of TDP-43.

### Hsp70 binds to the transient α-helical region of TDP-43 LCD

We next investigated the structural basis underlying the interaction between Hsp70 and TDP-43 LCD by using solution NMR spectroscopy. We prepared ^15^N-labeled TDP-43 LCD and titrated it by different concentrations of Hsp70. The 2D ^1^H-^15^N HSQC spectra showed a significant signal attenuation of a few residues upon Hsp70 titration, implying a direct binding between Hsp70 and the LCD of TDP-43 (Fig. 5a-b). Strikingly, the results pinpointed to the residues of amino acid (aa) 315-343 that counted for more than 50% of the signal attenuation (Fig. 5a-b). This region was previously identified to adopt a transient α-helical conformation, which is essential for mediating the LLPS and amyloid aggregation of TDP-43 (Conicella et al., 2016; Jiang et al., 2016; Conicella et al., 2020).

**Fig. 5:**
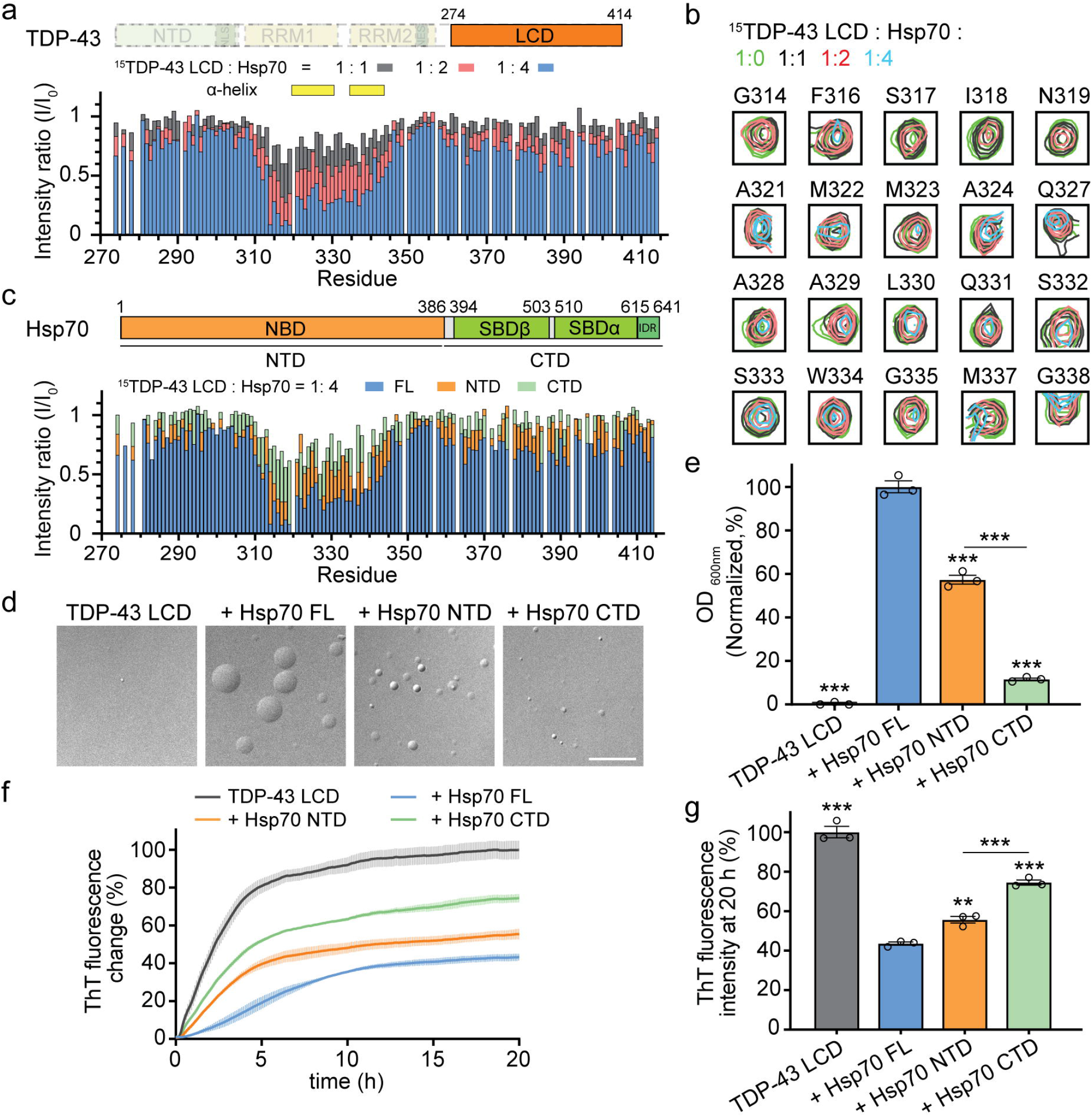
Structural characterization of the interaction between TDP-43 and Hsp70. **a** Residue-specific intensity changes of signals in the 2D ^1^H-^15^N HSQC spectra of ^15^N-labeled TDP-43 LCD (20 μM) in the presence of different concentrations of Hsp70 as indicated. The yellow blocks represent the previously identified transient □-helix region in TDP-43 LCD. **b** Representative residues with 2D ^1^H-^15^N HSQC intensity signal attenuation larger than 50% in (a, 1:4 ratio). **c** Schematic of the Hsp70 domains (upper). NBD, nucleotide-binding domain; SBD, substrate-binding domain (divided into SBDß and SBD); IDR, intrinsically disordered region. Residue-specific intensity changes of signals in the 2D ^1^H-^15^N HSQC spectra of ^15^N-labeled TDP-43 LCD (20 μM) with Hsp70 FL or truncations (80 μM) (lower). **d-e** Representative DIC images (d) and turbidity measurement (e) of TDP-43 LCD (50 μM) alone or mixed with Hsp70 FL or truncations (75 μM) in 50 mM NaCl, pH 6.0. **f-g** The ThT fluorescence assay (f) of TDP-43 LCD (20 μM) with Hsp70 FL or truncations (4 μM). The graph (g) shows ThT fluorescence intensity at 20 h in f. Mean ± SEM; n=3 in (e-g). Student’s *t* test (e and g); *\*\*p* < 0.01, and \*\*\**p* < 0.001. Scale bar, 10 μm in (d).

Hsp70 contains a N-terminal nucleotide-binding domain (NBD), a substrate-binding domain (SBD), and a C-terminal intrinsically disordered region (IDR) (Fig. 5c). To dissect which domain of the Hsp70 protein was responsible for interacting with TDP-43 LCD, we purified the truncated Hsp70-NTD (aa 1-385) and Hsp70-CTD (aa 386-641) proteins. The two Hsp70 truncations were titrated to ^15^N-labeled TDP-43 LCD, respectively. The 2D ^1^H-^15^N HSQC spectra revealed that similar to the FL Hsp70, titration of Hsp70-NTD induced a remarkable intensity attenuation especially within the residues of aa 315-343 that contained the transient α-helix region (Fig. 5c). In contrast, the signal attenuation caused by Hsp70-CTD was much weaker including in the α-helix region of TDP-43 LCD (Fig. 5c). Moreover, the NTD of Hsp70 exhibited a significantly stronger promotion of the LLPS of TDP-43 LCD than Hsp70-CTD (Fig. 5d-e). Also, Hsp70-NTD was more potent than Hsp70-CTD in preventing fibrillation of TDP-43 LCD (Fig. 5f-g). Together, our data indicated that Hsp70 directly interacted with the transient α-helix region of TDP-43 LCD mainly via the NTD, which stabilized TDP-43 in the liquid-like phase and prevented it from amyloid fibrillation.

### Hsp70 alleviates abnormal aggregation of ALS-associated TDP-43 mutant in cells

To demonstrate that the interaction between Hsp70 and TDP-43 underlay the recruitment and the anti-aggregation effect of Hsp70 in TDP-43 NBs, it would be ideal to test on TDP-43 mutants with disrupted Hsp70–TDP-43 interface. Toward this end, we prepared two TDP-43 LCD variants – a deletion of the residues of aa 313-319 (LCD Δ313-319), which was adjacent to the α-helix and exhibited a large signal attenuation upon Hsp70 titration, and an A326P mutation, which disrupted the transient α-helix of TDP-43 LCD (Conicella et al., 2016). Unfortunately, compared to WT, both variants showed severely impaired ability to phase separate *in vitro* (Fig. S5a-b) and were unable to form stress-induced NBs in cells (Fig. S5c-k), which excluded us from directly determining how the interaction and co-LLPS of Hsp70 with TDP-43 impacted on the dynamics, maturation and aggregation of TDP-43 in cells.

We then took an alternative approach to examine the chaperone activity of Hsp70 against pathological aggregation of TDP-43 NBs. An ALS/FTD kindred was recently reported to be associated with a mutation of TDP-43 at K181E (TDP-43-K181E), which was unable to bind to RNA and formed abundant aggregates in transfected cells (Chen et al., 2019). Indeed, cells expressing TDP-43-K181E spontaneously formed nuclear inclusions (NIs) in the absence of stress, of which a great portion showed large TDP-43 nuclear aggregates. Hsp70 was found to co-localize with both small and large TDP-43-K181E NIs, however, only the large ones were hyperphosphorylated (pTDP-43) (Fig. 6a-e). Of note, pTDP-43 at S409/410 was a histopathological hallmark of ALS (Neumann et al., 2009).

**Fig. 6:**
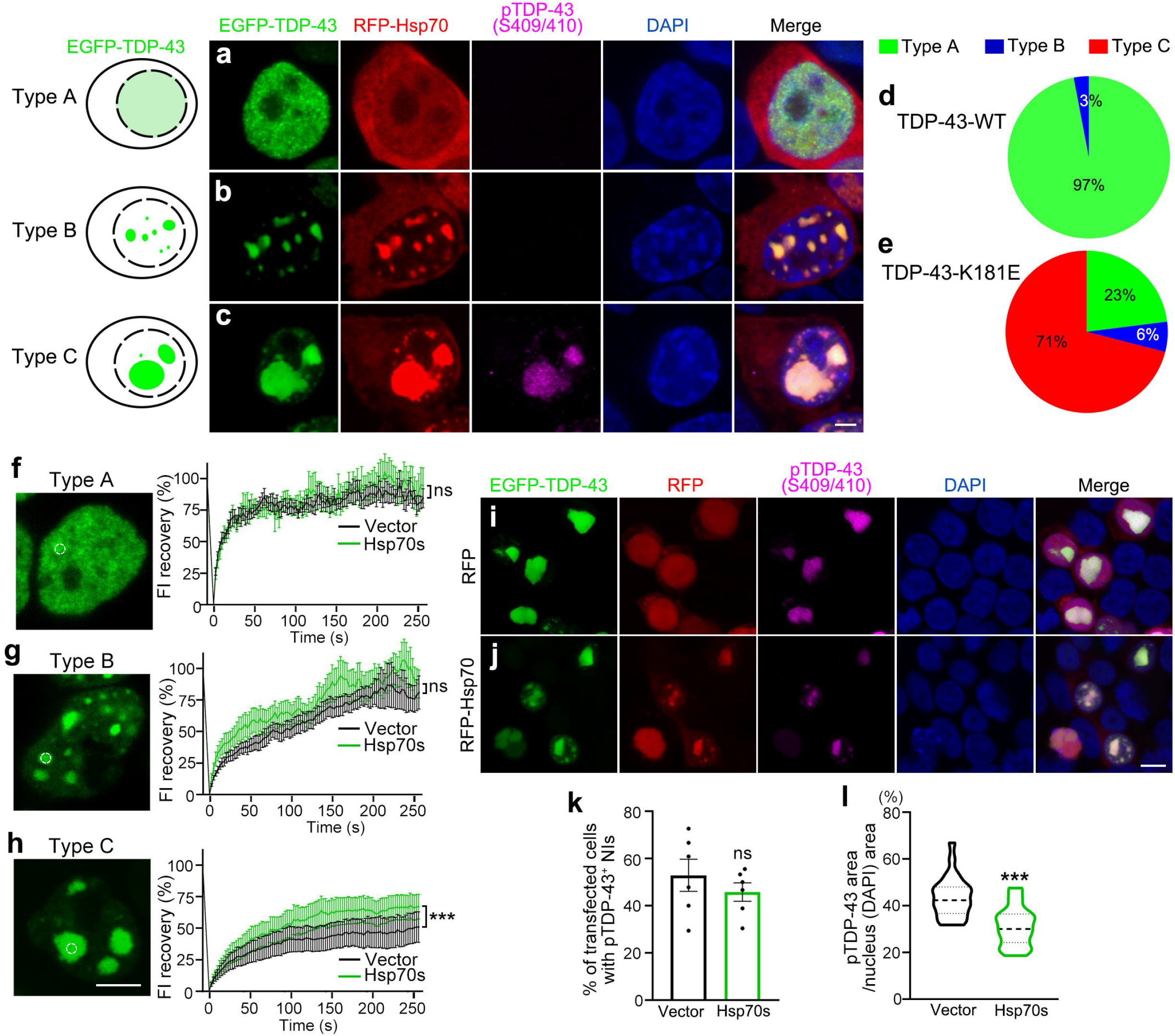
Upregulation of Hsp70 suppresses pathological aggregation of ALS-associated TDP-43-K181E mutation in the nucleus. **a-c** Representative illustrations and confocal images of three types of 293T cells when expressing WT or K181E EGFP-TDP-43: Type A, TDP-43 is diffused in the nucleus (a); Type B, TDP-43 forms small NBs or NIs (b); Type C, TDP-43 forms large NIs (c). RFP-Hsp70 is co-localized with TDP-43 in both Type B and Type C cells, but only Type C is immune-positive with anti-pTDP-43 (Ser409/410). **d-e** Classification and quantification of different TDP-43 morphology according to a-c in cells expressing TDP-43-WT (d) or TDP-43-K181E (e). **f-h** The FRAP assay evaluating the impact of OE of Hsp70s (*HSPA1A* and *HSPA8*) in living 293T cells on the dynamics of different types of EGFP-TDP-43-K181E nuclear inclusions. The dashed circles indicate the regions of the same size are photobleached in different types of TDP-43 nuclear inclusions. **i-j** Representative confocal images of 293T cells expressing EGFP-TDP-43 K181E and RFP (i) or RFP-Hsp70 (j), immunostained with pTDP-43 (Ser409/410) antibody and DAPI. **k-l** Quantifications of the percentage of transfected cells showing pTDP-43-positive nuclear inclusions (k) and the area of pTDP-43 (Ser409/410) nuclear inclusions normalized to the nuclear area (l). Mean ± SEM; n = ~100 cells in (d-e), n = 8 cells in (f-h), n = ~60 cells in (k) and n = 25 cells in (l). Two-way ANOVA in (f-h), Student’s *t* test in (k-l); \*\*\**p* < 0.001; ns, not significant. Scale bars, 5 μm in (a-c and f-h) and 10 μm (i and j).

Further, the FRAP assay revealed that, compared to diffused nuclear TDP-43-K181E or small TDP-43-K181E NIs, large TDP-43-K181E NIs recovered the most slowly (Fig. 6f-h). More importantly, we showed that OE of Hsp70s (together of *HSPA1A* and *HSPA8*) not only markedly increased the dynamics of large TDP-43-K181E NIs (Fig. 6h), but also significantly reduced the pTDP-43 levels of the K181E mutation in the nucleus (Fig. 6i-l). Collectively, these data indicated that Hsp70 could chaperone TDP-43 NBs in dynamic, liquid-like phase, which prevented them from forming pathological aggregation in cells under stressed or diseased conditions.

## DISCUSSION

Cellular stress triggers TDP-43 to undergo LLPS and condense into cytoprotective NBs. In a recent study (Wang et al., 2020), we have demonstrated that WT TDP-43 does not spontaneously phase separate into subnuclear compartments in normal cells, as the high content of RNAs in the nucleoplasm suppresses TDP-43 LLPS. In response to stress, the expression of lncRNA *NEAT1* is upregulated, which binds to TDP-43 preferentially via the RRM1 and promotes the assembly of TDP-43 NBs. In the current work, we continue to investigate the mechanisms regulating stress-induced TDP-43 NBs. We find that Hsp70 can directly bind to the highly aggregation-prone α-helix region in the LCD of TDP-43. Thus, although both *NEAT1* and Hsp70 appear to promote the assembly of TDP-43 NBs, they recognize different regions of TDP-43, act through different mechanisms, and play different roles in the regulation of TDP-43 NBs.

Previous studies show that both the LCD and ΔLCD (which contains the RRMs) of TDP-43 are capable of LLPS (Schmidt and Rohatgi, 2016; Duan et al., 2019; Wang et al., 2020), however, the LCD plays a predominant role in mediating pathological amyloid aggregation of TDP-43 (Johnson et al., 2009). While *NEAT1* preferentially binds to the RRM1 and provides a “nucleation core” that triggers the assembly of TDP-43 NBs (Wang et al., 2020), Hsp70 binds to the aggregation-prone region of TDP-43, which not only co-phase separates with TDP-43 but also potently suppresses the liquid-to-solid phase transition of TDP-43 NBs, keeping them in the highly dynamic, liquid-like phase. We notice that a recently posted preprint reports that disrupting the RNA binding of TDP-43 by acetylation or mutations in the RRMs causes TDP-43 to form intranuclear liquid spherical annuli, of which Hsp70 was identified as a key component (Yu et al., 2020). The findings are in line with our conclusion that Hsp70 plays a crucial role in chaperoning TDP-43 NBs.

Hsp70 is a master chaperone in protein quality control, and it serves as a central physical node for binding other chaperones, client protein and co-chaperones (Kampinga and Craig, 2010). The NBD of Hsp70 binds to only not nucleotides but also other chaperones such as Hsp40 and Hsp110 and co-chaperones such as Bag1 (Sondermann et al., 2001; Jiang et al., 2007; Polier et al., 2008). Nucleotide-binding and ATP hydrolysis in the NBD mediate the conformational rearrangement, which controls the binding and release of clients and other chaperones (Rosenzweig et al., 2019). Intriguingly, our study reveals that the NBD mediates the binding of Hsp70 to TDP-43. It will be interesting to explore whether and how ATP/ADP and the binding partners such as Hsp40 and other co-chaperones are involved in the regulatory role of Hsp70 in the assembly and maintenance of TDP-43 NBs.

Earlier studies report that upregulation of Hsp70 can suppress the age-dependent degeneration of the fly eye caused by OE of either WT or A315T mutant TDP-43 (Estes et al., 2011), and Hsp70 prevents aggregate accumulation of the 25-kDa C-terminal fragment of TDP-43 (Kitamura et al., 2018). In this study, we find that although OE of Hsp70 is insufficient to alter the assembly/disassembly, maturation or cytotoxicity of WT TDP-43 NBs, it potently delays maturation of the ALS-causing mutant TDP-43-K181E and significantly decreases the pTDP-43 levels without affecting the number of cells forming TDP-43-K181E NIs. These data suggest that Hsp70 is less likely a key factor triggering the assembly of TDP-43 NBs. Rather, it plays a major role in promoting the dynamics of TDP-43 NBs and maintaining them in the liquid-like, non-pathological state. As such, the beneficial effect of OE of Hsp70 is more prominent with the disease-causing TDP-43-K181E mutation. Intriguingly, expression of Hsp70 is decreased in ALS patients associated with TDP-43 pathology (Chen et al., 2016), and OE of TDP-43 in both fly and mouse motor neurons leads to reduced expression of Hsp70 (Coyne et al., 2017). Therefore, dysregulation of Hsp70 in diseased conditions may contribute to and further worsen liquid-to-solid transition and aggregation of TDP-43 in ALS and other related diseases.

## MATERIALS AND METHODS

### Plasmid construction

The pcDNA3.1-TDP-43-HA, pCMV-myc-TDP-43 and pCAG-EGFP-TDP-43 plasmids were used as previously described (Sun et al., 2018; Wang et al., 2020). The pcDNA3.1-TDP-43-HA and pCAG-EGFP-TDP-43 plasmids were used as the templates to generate the following plasmids. The pcDNA3.1-TDP-43-A326P-HA, pcDNA3.1-TDP-43-Δ(313-319)-HA and pCAG-EGFP-TDP-43-K181E plasmids were generated by site-directed mutagenesis using the Fast Mutagenesis Kit II (Vazyme). To generate the pCAG-RFP-Hsp70 and pCAG-RFP-Hsc70 plasmids, the Hsp70, Hsc70 and RFP coding sequence were amplified from pET28a-His-Hsp70 (a gift from Dr. P. Chen), pET28a-His-SUMO-Hsc70 (a gift from Dr. L. He) and pCAG-RFP, respectively, and then subcloned into the pCAG plasmid using the ClonExpress MultiS One Step Cloning Kit (Vazyme). The pcDNA3.1-Hsp70-HA, pcDNA3.1-Hsc70-HA, pCMV-myc-Hsp70 and pCMV-myc-Hsc70 plasmids were generated using the same way.

For *E. coli* expression, genes of TDP-43 ΔLCD (aa 1-273) and TDP-43 LCD (aa 274-414) plasmids were used as previously described (Duan et al., 2019). FL Hsp70 were cloned into pET28a with a N-terminal His-tag. TDP-43 LCD variants including TDP-43 LCD A326P, Δ313-319 were generated from pET28a TDP-43 LCD wild-type (WT). The Hsp70 truncations including Hsp70-NTD (aa 1-385) and Hsp70-CTD (aa 386-641) were generated from FL Hsp70. TDP-43 was cloned into pET9d with an TEV protease cleavable MBP-His tag at C-terminus. Hsc70 was subcloned into pET28a vector with a N-terminal His-SUMO tag. All constructs were verified by sequencing and the primers used for PCR to generate the expression plasmids are summarized as following:

pcDNA3.1-TDP-43-A326P-HA and pET28a-His-TDP-43-LCD-A326P:
5’ - CATGATGGCTGCCCCGCAG -3’
5’ - TAGTGCTGCCTGCGGGGC -3’;
pcDNA3.1-TDP-43-Δ(313-319)-HA and pET28a-His-TDP-43-LCD-Δ(313-319):
5’ - GATGAACCCAGCCATGATGGCTGCC -3’
5’- ATCATGGCTGGGTTCATCCCACCACCCATATTAC -3’
pCAG-GFP-TDP-43-K181E:
5’ - TTCTGAGCAAAGCCAAGATGAGCCTTTGAGAA -3’
5’ - CTTGGCTTTGCTCAGAATTAGGAAGTTTGCAGTCACACC -3’
pCAG-RFP-Hsp70:
5’ - CATCATTTTGGCAAAGAATTCGCCACCATGGCCTCCTCCGAGGACGTC -3’
5’ - CTTTGGCCATGCTTCCGCCGGCGCCGGT -3’
5’- CGGCGGAAGCATGGCCAAAGCCGCGGCG -3’
5’ - GCTCCCCGGGGGTACCTCGAGCTAATCTACCTCCTCAATGGTGGG -3’ pCAG-RFP-Hsc70:
5’ - CATCATTTTGGCAAAGAATTCGCCACCATGGCCTCCTCCGAGGACGTC -3’
5’- AGGTCCCTTGGACATGCTTCCGCCGGCGCCGGT -3’
5’ - GAAGCATGTCCAAGGGACCTGCAGTT -3’
5’- GCTCCCCGGGGGTACCTCGAGTTAATCAACCTCTTCAATGGTGGG -3’ pcDNA3.1-Hsp70-HA:
5’-CGTTTAAACGGGCCCTCTAGAGCCACCATGGCCAAAGCCGCGGCGATC -3’
5’-ATATCCAGCACAGTGGCGGCCGCTTAAGCGTAGTCTGGGACGTCGTATGGGTAATCTACCTCCTCAATGGTGGGGCC -3’
pcDNA3.1-Hsc70-HA:
5’- CGTTTAAACGGGCCCTCTAGAGCCACCATGTCCAAGGGACCTGCAGTTG -3’
5’- ATATCCAGCACAGTGGCGGCCGCTTAAGCGTAGTCTGGGACGTCGTATGGGTAATCAACCTCTTCAATGGTGG -3’
pCMV-myc-Hsp70:
5’- ATGGCCATGGAGGCCGAATTCGCCACCATGGCCAAAGCCGCGGCGATC -3’
5’- CCGCGGCCGCGGTACCTCGAGCTAATCTACCTCCTCAATGGTGGG -3’ pCMV-myc-Hsc70:
5’- ATGGCCATGGAGGCCGAATTCGCCACCATGTCCAAGGGACCTGCAGTTG -3’
5’- CCGCGGCCGCGGTACCTCGAGCTATTAATCAACCTCTTCAATGGTGGG -3’
pET28a-Hsp70-NTD:
5’- CAGCAAATGGGTCGCGGATCCATGGCCAAAGCCGCGGCG -3’
5’ - GTGTGGTGGTGGTGCTCGAGTTACTCGGACTTGTCCCCCA -3’
pET28a-Hsp70-CTD:
5’ - CAGCAAATGGGTCGCGGATCCAACGTGCAGGACCTGCTGC -3’
5’ - GTGGTGGTGGTGGTGCTCGAGCTAATCTACCTCCTCAATGGTGGG -3’ pET9d-TDP-43-TEV-MBP-His:
5’ - CTTTAAGAAGGAGATATACCATGTCTGAATATATTCGGGTAACCG -3’
5’-CCGCCTCCCTGAAAATAAAGATTCTCGCTTCCGCCCATTCCCCAGCCAGAAGA CTTA -3’
5’- CTTTATTTTCAGGGAGGCGGAAGCGGCGGAAGCATGAAAATCGAAGAAGGTAAACG -3’
5’ - TGCCATAGCTACTGCTGCTGTTAATGATGATGATGATGGTGCATA -3’

### Protein expression and purification

For protein purification, TDP-43 LCD WT and its variants were overexpressed in BL21(DE3) *E. coli* cells by adding 1 mM IPTG with incubation at 37 °C for 12 h. Cells were harvested and lysed in 50 mM Tris-HCl, pH 8.0, and 100 mM NaCl. Cell pellet was collected by centrifugation (16,000 rpm, 4 °C, 1 h), and then was resuspended into the denatured buffer containing 50 mM Tris-HCl, pH 8.0, 6 M guanidine hydrochloride with sonication. The resuspended protein was loaded onto a Ni column (GE Healthcare, USA) after filtration. Protein was then eluted by the denatured elution buffer containing 50 mM Tris-HCl, pH 8.0, 6 M guanidine hydrochloride and 100 mM imidazole. The eluted protein was desalted into storage buffer (20 mM NaPhosphate, pH 7.0 and 8 M urea), and further concentrated into over 30 mg/ml proteins, and flash frozen and kept at −80 °C for storage. For further experiments, the protein was desalted into the buffer containing 20 mM MES, pH 6.0.

TDP-43 ΔLCD was overexpressed in Bl21(DE3) *E. coli* cells and Hsp70s were overexpressed in BL21(DE3) codon plus *E. coli* cells. Protein expression was induced by adding 0.5 mM IPTG at 22 °C for 12 h. For protein purification, His-TDP-43 ΔLCD were lysed in lysis buffer consisting of 50 mM Tris-HCl, pH 7.5, 500 mM NaCl, 20 mM imidazole, 4 mM β-mercaptoethanol, and 2 mM PMSF. RNase A (0.1mg/ml) was added into lysis buffer to remove TDP-43 ΔLCD binding RNA. Proteins were loaded on Ni column and further eluted with Ni elution buffer containing 50 mM Tris-HCl, pH 7.5, 500 mM NaCl, 250 mM imidazole and 4 mM β-mercaptoethanol. Eluted proteins His-TDP-43 ΔLCD was further desalted into storage buffer containing 50 mM Tris-HCl, pH 7.5, 500 mM NaCl and 2 mM DTT.

As for purification of Hsp70s, His-SUMO-Hsc70, His-Hsp70 WT and its variants were lysed in lysis buffer and eluted with Ni elution buffer, and then eluted proteins including His-Hsp70 WT and its variants were fractionated via Superdex 200 16/600 column in 50 mM Tris-HCl, pH 7.5, 100 mM NaCl and 2 mM DTT. As for Hsc70, His-Ulp1 was used to remove His-SUMO tag before gel-filtration.

FL TDP-43 MBP was overexpressed in BL21(DE3) PlySs *E. coli* cells with 1 mM IPTG at 16 °C for 16 h. Cells were harvested and lysed in 50 mM Tris-HCl, pH 7.5, 1 M NaCl, 2 mM DTT, 10% glycerol, 1 mM EDTA and 2 mM PMSF. After removing cell pellets by centrifugation, protein was loaded onto MBP Trap HP column and then eluted with 50 mM Tris-HCl, pH 7.5, 1 M NaCl, 2 mM DTT, 10% glycerol, and 10 mM maltose. Eluted protein was purified over the gel filtration chromatography (Superdex 200 16/300; GE Healthcare) equilibrated with storage buffer (50 mM Tris-HCl, pH 7.5, 300 mM NaCl, 2 mM DTT).

### Fluorescent labeling of the proteins

For labeling proteins with active thiol groups, proteins were desalted into reaction buffer (50 mM Tris-HCl, pH 7.5, 500 mM NaCl and 4 mM Tris (2-Carboxyethyl) Phosphine (TCEP) (Invitrogen, T2556)) for removing DTT in storage buffer. The proteins were then incubated with a 5-fold fluorescent dye (molar ratio), including Alexa 488 C_5_-malemide (Invitrogen, A10254) for FL TDP-43 MBP, Alexa 555 C2-malemide (Invitrogen, A20346) for TDP-43 ΔLCD, and Alexa 647 C_2_-malemide (Invitrogen, A20347) for Hsp70 and Hsc70. The labeling reaction was performed for over 1 h at room temperature (RT). The labeled proteins were further purified using the Superdex 200 10/300 columns (GE Healthcare, USA). As for TDP-43 LCD, the protein stored in denatured buffer was desalted with HPLC (Agilent) and freeze-dried by the FreeZone lyophilizer (Thermo Fisher). The protein powder was dissolved into 50 mM NaPhosphate (pH 7.0). Then, the protein solution was incubated with 10-fold OregonGreen488 (Invitrogen, O6149) (molar ratio) at 37 °C for 1 h. The reaction was then mixed with 20-fold volume of denature buffer containing 50 mM Tris-HCl, pH 8.0 and 8 M urea, and further purified via Superdex 75 16/600 column in 20 mM MES, pH 6.0. For further LLPS assay and confocal imaging, the unlabeled protein was mixed with labeled one at the molar ratio of 49:1 (unlabeled: labeled).

### *In vitro* LLPS assay

For co-LLPS experiments, 50 μM FL TDP-43 MBP were mixed with 10 μM Hsp70 or 10 μM Hsc70 under 50 mM Tris-HCl, pH 7.5, 150 mM NaCl, 2 mM DTT and 15% Dextran 70. Protein phase separation was initiated with the addition of Dextran 70. As for co-LLPS between Hsp70 and TDP-43 LCD or ΔLCD, Hsp70 was added at last step to achieve final conditions as indicated. To compare phase separation ability between TDP-43 LCD WT and its variants, 4 M NaCl was added to 20 μL proteins to achieve final conditions as indicated.

### Turbidity measurement

Turbidity of different samples were measured based on the optical absorption at 600 nm. The measurements were recorded on a Varioskan Flash spectral scanning multimode reader (Thermo Fisher) using a flat bottom 384-well plates (20 μL pre well, Corning).

### DIC and fluorescent imaging for phase separated protein samples

LLPS samples were loaded onto glass slide with coverslip. DIC and confocal images were acquired on a Leica TCS SP8 microscope with a 100 × objective (oil immersion) at RT.

### Nuclear magnetic resonance (NMR)

Backbone assignment of TDP-43 LCD was accomplished according to the previous publication (Conicella et al., 2016). All NMR titration experiments were performed at 298 K on a Bruker 900 MHz spectrometer equipped with cryogeni probe in an NMR buffer of 20 mM MES (pH 6.0), 150 mM NaCl, 10% glycerol and 20% D_2_O. Here, we increased salt concentration and utilized glycerol to avoid LLPS of TDP-43 LCD with Hsp70. Each NMR sample was made with a volume of 500 μL, containing ^15^N-TDP-43 LCD (20 μM) desalted from denature buffer freshly with/without Hsp70 WT and its variants as indicated. Bruker standard pulse sequence (hsqcetfpf3gpsi) was used to collect the 2D ^1^H-^15^N HSQC spectrum with 16 scans. And 2048 × 160 complex points were used for ^1^H (14 ppm) and ^15^N (21 ppm) dimension, respectively. All NMR data were processed by NMRpipe and analysed by SPARK (Delaglio et al., 1995; Lee et al., 2015).

### ThT fluorescence kinetic assay

ThT fluorescence kinetic assay was performed in the ThT assay buffer containing 20 mM MES, pH 6.0, 150 mM NaCl, 4 mM DTT, and 0.05% NaN3 with 20 μM TDP-43 LCD and Hsp70 or its variants, respectively. The mixture was added into a 384-well plate (Corning) with 50 mM ThT. The ThT fluorescence was monitored by a Varioskan Flash spectral scanning multimode reader (Thermo Fisher Scientific) with excitation at 440 nm and emission at 485 nm at 37 °C and the plate was shaken at 900 rpm. The morphology of TDP-43 LCD fibril was visualized by TEM.

### Negative-staining transmission electron microscopy (TEM)

Samples were incubated on carbon-coated grids for 1 min and washed with ddH_2_0 for twice after staining with 8 μL uranyl acetate (2%, v/v) for 45 s. The grids were further assessed by using Tecnai G2 Spirit transmission electron microscope with 120 kV voltage. The TEM images were obtained by a 4000 × 4000 charge-coupled device camera (BM-Eagle, FEI Tecnai).

### Cell cultures and transfection

HEK293T and HeLa cells were cultured in Dulbecco’s Modified Eagle Medium (DMEM, Gibco) supplemented with 10% (v/v) fetal bovine serum (FBS, VISTECH) and 1% penicillin/streptomycin at 37 °C in 5% CO2. Transient transfection was performed using PolyJetTM (SignaGen) in DMEM. Cells were transfected for at least 24 h before the subsequent drug treatments or examinations. For the knockdown experiment, the siRNA (Genepharma) was transfected into the HeLa cells using the LipofectamineTM RNAiMax Transfection Reagent (Invitrogen) according to the manufacturer’s instruction. The siRNA was incubated for 48-72 h before cells were harvested. The siRNA oligos used in this study are listed below:

si-Ctrl: 5’- UUCUCCGAACGUGUCACGUTT -3’;
si-Hsp70: 5’- CCAAGCAGACGCAGAUCUUTT -3’;
si-Hsc70: 5’- GCUGUUGUCCAGUCUGAUATT -3’

### Arsenite treatment and washout assay

HeLa cells were grown on the coverslips in the 24-well plate and transfected with the indicated plasmids or siRNA. Cells were then treated with 250 μM NaAsO2 or PBS for 30min, 1h or 2h at indicated. For arsenite washout assay, the arsenite-containing medium was removed and washed in PBS, the cells were then incubated in fresh culture medium for the indicated time.

### Immunocytochemistry and confocal imaging

HeLa or HEK293T cells were grown on coverslips in the 24-well plate and HEK293T cells were pre-coated with PLL (Sigma) in a 24-well plate before transcription and treatment. The cells were then fixed in 4% paraformaldehyde in PBS for 30 min at RT, permeabilized in 0.5% TritonX-100 (Sigma) in PBS for 30 min and blocked with 3% goat serum in PBST (0.3% goat serum in PBS for pTDP-43) for 1 h at RT. The primary and secondary antibodies were then incubated in the blocking buffer at 4 °C overnight, or at RT for 1-2 h. After washing for 3 times with PBST (PBS for pTDP-43), cells were mounted on glass slides using the VECTASHIELD Antifade Mounting Medium with DAPI (Vector Laboratories). Fluorescent images were taken using the Leica TCS SP8 or Light Sheet confocal microscopy system using 63 × or 100 × oil objective (NA = 1.4).

### FRAP assay

FRAP assay was performed using the FRAP module of the Leica SP8 or Light Sheet confocal microscopy system using 100 × oil objective (NA= 1.4). For living cells, the EGFP-TDP-43 NB was bleached using a 488 nm laser at 100% laser power for twice. Bleaching was focused on a 1 μm diameter region of interest. After photobleaching, time-lapse images were captured every 8 s for about 5 min. As for *in vitro* FRAP, the assay was performed in similar way. An aliquot of 20 μL LLPS sample was applied to a glass bottom dish (40 mm × 40 mm, 0.2 mm at thinnest bottom) (Nest, 80100). Bleaching was also focused on the same size at the droplets with similar diameter in the same group. After photobleaching, images were continuously captured (1 image/2.58s). As for analysis of FRAP data, the fluorescent intensity (I_t_^m^) recorded on the bleached region in each time point (t) were normalized to fluorescent intensity (I_t_^c^) of nearby unbleached region, with the formula: It=(I_t_^m^/I_0_^m^)/(I_t_^c^/I_0_^c^). Fluorescence recovery fraction for bleached intensity was further calculated with the formula: (I_t_ - I_min_)/ (I_0_ - I_min_). I_min_ is the unbleached fraction after photobleaching. ImageJ and GraphPad Prism are used to measure and analyze the FRAP data.

### RNA extraction and real-time quantitative PCR (qPCR)

For (qPCR), total RNA was isolated from HeLa cells using TRIzol (Invitrogen) according to the manufacturer’s instruction. After DNase (Promega) treatment, the reverse transcription reactions were performed using reverse Transcriptase M-MLV (RNase H-) (TaKaRa). The cDNA was then used for real-time qPCR using the SYBR Green qPCR Master Mix (Bi-make) with the QuantStudio 6 Flex Real-Time PCR system (Life Technologies). The mRNA levels of β-actin were used as an internal control to normalize the mRNA levels of Hsp70s. The qPCR primers used in this study are listed below:

below:

*HSPA8:*
5’ - TTGGAGTGGTTCGGTTTCCC -3’
5’ - TATTGGAGCCAGGCCTACAC -3’;
*HSPA9:*
5’ - CTTGTTTCAAGGCGGGATTATGC -3’
5’ - GCAGGAGTTGGTAGTACCCAAA -3’;
*HSPA5:*
5’ - CATCACGCCGTCCTATGTCG -3’
5’- CGTCAAAGACCGTGTTCTCG -3’;
*HSPA1A:*
AGCTGGAGCAGGTGTGTAAC
CAGCAATCTTGGAAAGGCCC;
*HSPA1B:*
5’ - TCTGGGTCAGGCCCTACCATT -3’
5’- AGCAGCAAAGTCCTTGAGTCC -3’;
*HSPA2:*
5’- AGATCGACTCGCTCTACGAGG -3’
5’- CGAAAGAGGTCGGCATTGAG -3’;
*HSPA1L:*
5’ - TTACCGTGCCAGCCTATTTCA -3’
5’- AGCACATTAAGTCCAGCAATCA -3’;
*hβ-actin:*
5’- GTTACAGGAAGTCCCTTGCCATCC -3’
5’ - CACCTCCCCTGTGTGGACTTGGG -3’;

### Cell viability assay

Transfected HeLa cells were seeded in 96-well plates at the density of 10^4^ cells/well and cultured in 100 μL of culture medium. After 48 h transfection, cells were treated with arsenite (100 μM) in different time points, and then cell viability was examined using the Cell Counting Kit-8 (CCK-8) (Dojindo), according to the manufacturer’s instructions. Briefly, 10 μL of the CCK-8 solution was added to each well and incubated at 37 °C for 2.5 h. Finally, the absorbance at 450 nm was measured with a Synergy2 microplate reader (BioTek Instruments).

### Antibodies

The following antibodies were used for Western blotting and immunocytochemistry assays: mouse anti-G3BP (BD Biosciences, 611127), rabbit anti-HA (CST, C29F4), rabbit anti-c-Myc (Sigma, c3956), mouse anti-pTDP-43 (Ser409/410, CosmoBio, CAC-TIP-PTD-M01), mouse anti-Hsp70 (StressMarq Biosciences Inc, SMC-100A/B), mouse anti-GAPDH (Proteintech, 60004-1). HRP conjugated secondary antibodies: goat anti-mouse (Sigma, A4416), goat anti-rabbit (Sigma, A9169). Fluorescent secondary antibodies: goat anti-rabbit-Alexa Flour 488 (Life Technologies, A11034), goat anti-mouse-Alexa Flour 568 (Life Technologies, A11031), goat anti-mouse-Alexa Flour Cy5 (Life Technologies, A10524).

### Statistical analysis

The statistical significance in this study is determined by the two-way ANOVA with Bonferroni’s post-hoc test or the unpaired, two-tailed Student’s *t*-test at \**p* < 0.05, \*\**p* < 0.01, and \*\*\**p* < 0.001.

## ACKNOWLEDGMENTS

We would like to thank the staffs in the National Center for Protein Science, Shanghai, for their assistance on NMR data collection. This work was supported by grants from the National Natural Science Foundation of China (NSFC) (91853113), the Science and Technology Commission of Shanghai Municipality (STCSM) (201409003300), the NSFC (81671254, 31872716 and 31970697), the STCSM (18JC1420500, 20490712600, 20XD1425000 and 2019SHZDZX02), and the “Eastern Scholar” project supported by Shanghai Municipal Education Commission.

## AUTHOR CONTRIBUTIONS

Y.F. and C.L. conceived the research; J.G., C.W., R.H., Y.F. and C.L. designed the project. J.G., C.W., R.H., Y.L., Y.S., Y.S. and Q.W. performed the experiments, J.G., C.W., R.H. and Q. W. contributed important new reagents; J.G., C.W., R.H., Y.F. and C.L. analyzed the data and interpreted the results; J.G., C.W. and R.H. prepared the figures; and D.L., Y.F. and C.L. wrote the manuscript. All authors read and approved the final manuscript.

