## Supplemental Information for "Hsp70 chaperons TDP-43 in dynamic, liquid-like phase and prevents it from amyloid aggregation"

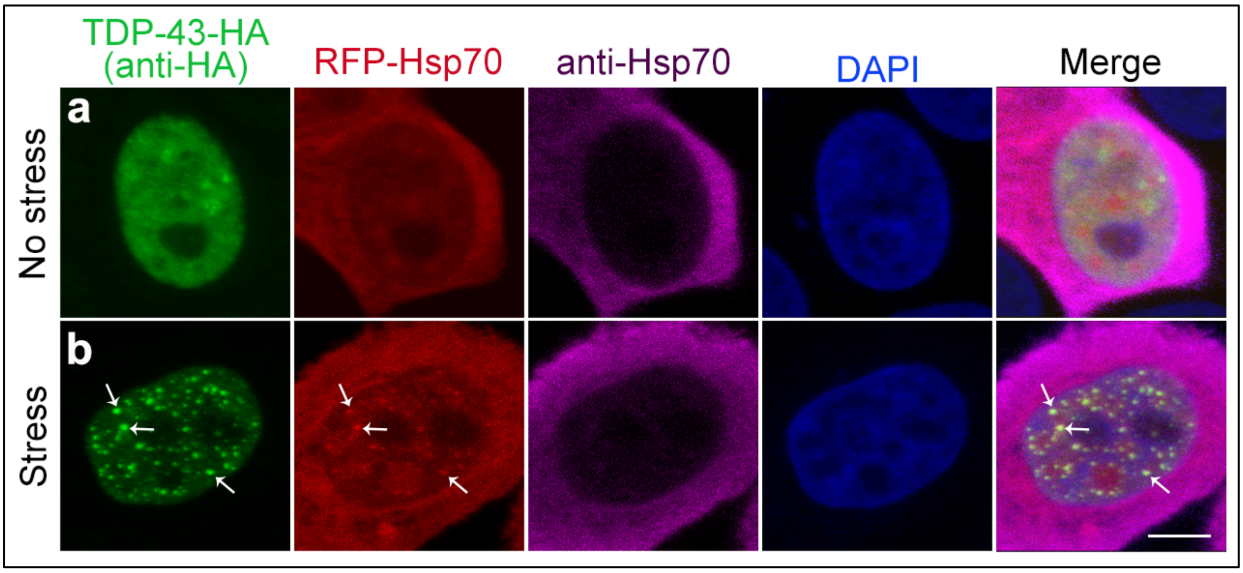


**Fig. S1: The anti-Hsp70 antibody fails to immunostain Hsp70 protein within TDP-43 NBs.**

**a-b** Representative images of HeLa cells co-expressing TDP-43-HA and RFP-Hsp70 in the absence (a) or presence (b) of arsenite (250 μM, 30 min). DAPI, nuclear labeling; anti-HA for TDP-43-HA; anti-Hsp70 for Hsp70. Scale bar, 5 μm. Arrows, co-localization of RFP-Hsp70 with TDP-43 NBs, which show no or minimal anti-Hsp70 signal.

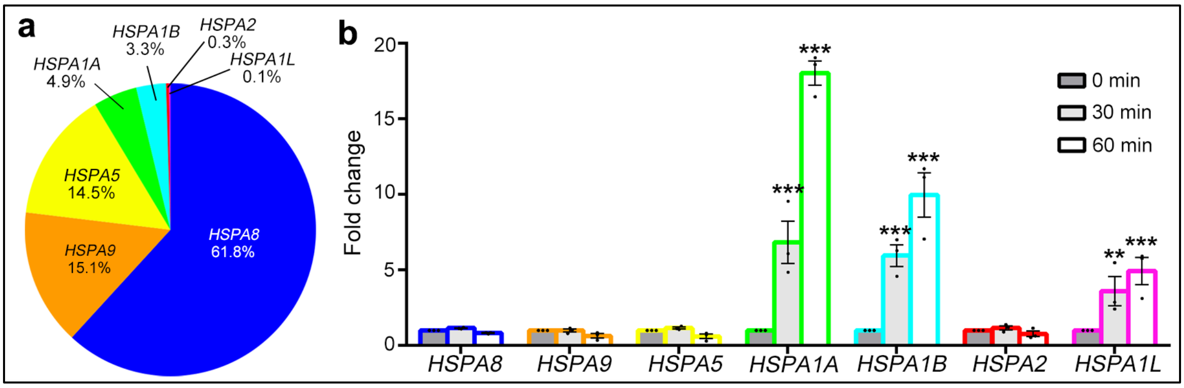


**Fig. S2: Among the Hsp70 family genes, *HSPA8* is expressed the most while *HSPA1A* responds the most robust under stress in HeLa cells.**

**a** Relative mRNA expression levels of different genes of the Hsp70 family are determined by qPCR in HeLa cells under normal condition. **b** The fold change of the mRNA levels of different genes of the Hsp70 family upon arsenite treatment (250 μM) for 0 min, 30 min or 60 min. All RNA levels were normalized to that of β-actin. Mean ± SEM; n = 3 in; One-way ANOVA; **$p < 0.01$; ***$p < 0.001$; ns, not significant.

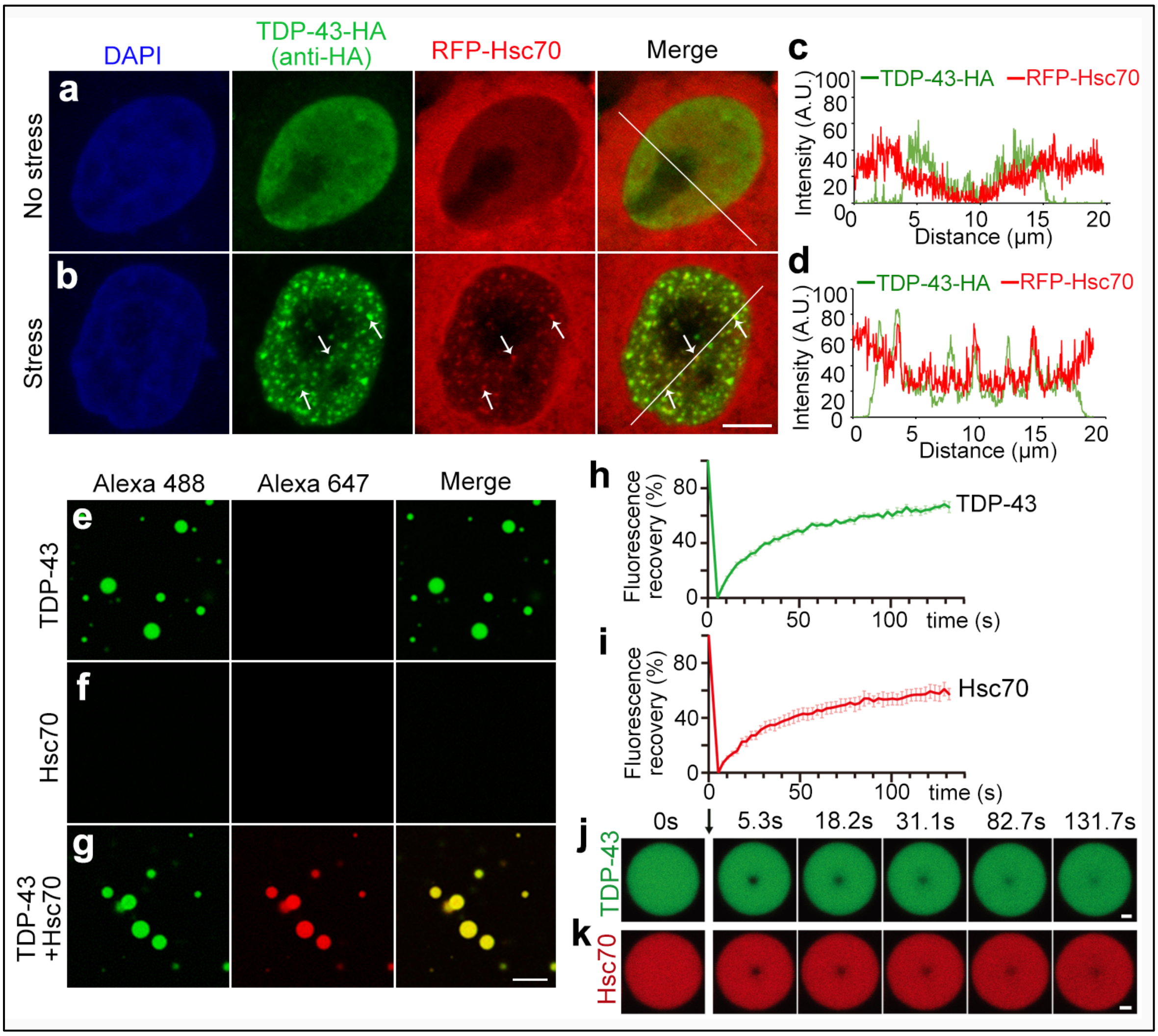


**Fig. S3: Hsc70 (encoded by *HSPA8*) co-localizes with TDP-43 NBs in stressed cells and co-phase separates with TDP-43 *in vitro*.**

**a-d** Representative images (a-b) and the line intensity analysis (c-d) of HeLa cells expressing RFP-Hsc70 together with TDP-43-HA in the absence or presence of cellular stress (250 μM of arsenite, 30 min). DAPI, nuclear labeling; anti-HA for TDP-43-HA; arrows, co-localization of RFP-Hsc70 with TDP-43 NBs. **e-g** Representative confocal images showing the *in vitro* LLPS of TDP-43-MBP (Alexa Fluor 488, green) alone (e), Hsc70 (Alexa Fluor 647, red) alone (f), and mix of them two (g). The concentration of each component in the *in vitro* LLPS assay: 50 mM Tris-HCl, pH 7.5, 150 mM NaCl, 15% Dextran 70, 50 μM TDP-43-MBP and 10 μM Hsc70. **h-k** The FRAP analyses (h-i) and images of representative droplets (j-k) of TDP-43 (h, j, green) and Hsc70 (i, k, red) in the co-phase separated droplets in g. The black arrow indicates the photobleaching. Scale bars, 5 μm in (a-b), 10 μm in (e-g) and 1 μm in (j-k). Means ± SEM, n = 5 in (h-i).

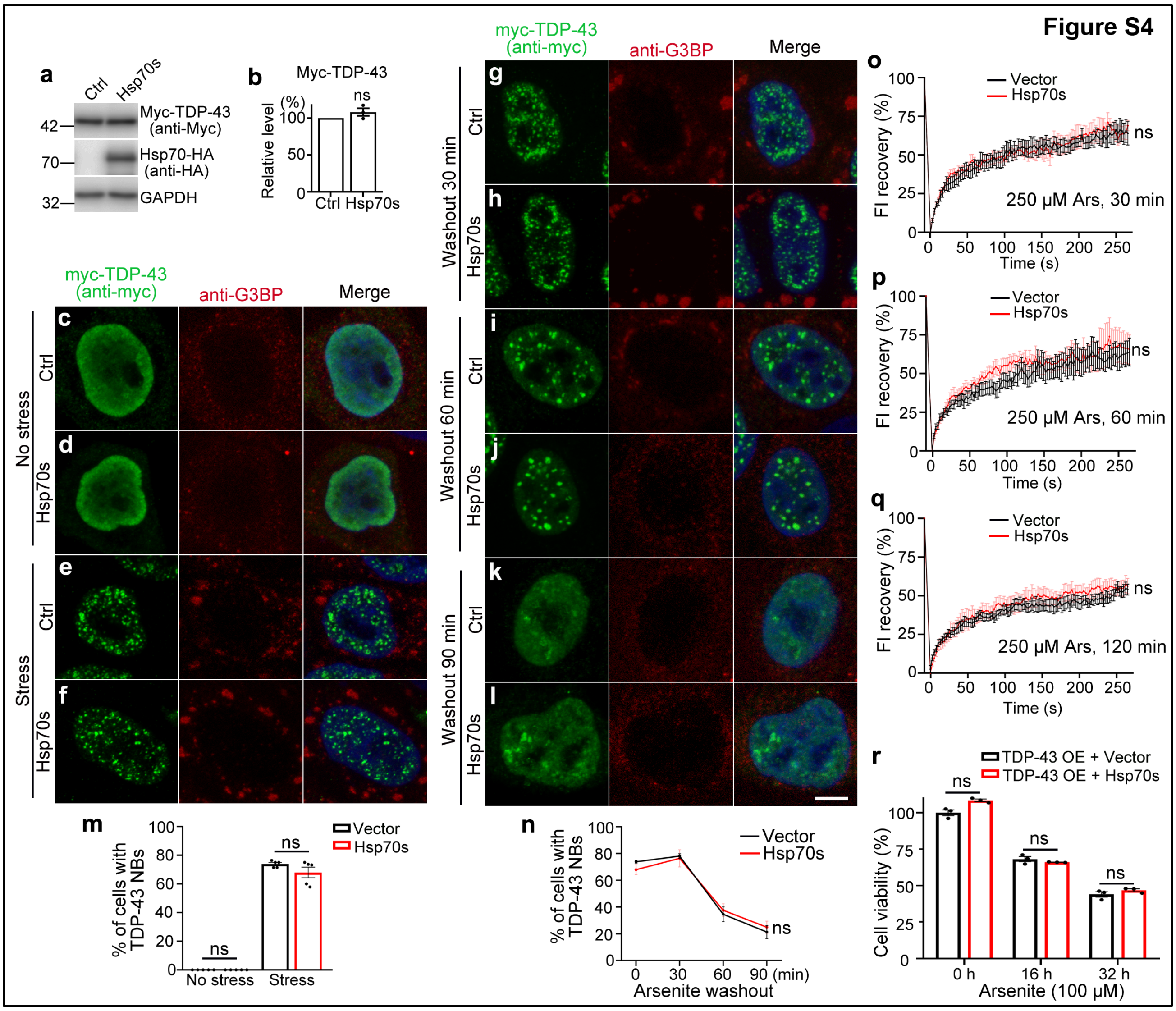


**Fig. S4: Hsp70s OE has no obvious effect on TDP-43 NBs.**

**a-b** Representative images (a) and quantification (b) of Western blot analysis confirming the expression of transiently transfected HA-Hsp70 and HA-Hsc70 (indicated as Hsp70s) and myc-TDP-43. **c-f** Representative confocal images of HeLa cells expressing myc-TDP-43 with vector or Hsp70s, treated with PBS (c-d) or 250 μM of arsenite for 30 min (e-f). **g-l** Representative images of the washout assay at 30 min (g-h), 60 min (i-j) and 90 min (k-l) after replacing the arsenite medium with normal medium. Merge with DAPI (for nucleus) is shown. anti-G3BP for SGs. **m** The percentage of cells forming stress-induced TDP-43 NBs in e-f). **n** The percentage of cells showing TDP-43 NBs at indicated time points after washout in e-l. **o-q** Fluorescent intensity recovery curves of EGFP-TDP-43 NBs in the FRAP assay. Cells expressing EGFP-TDP-43 with vector or Hsp70s treated with arsenite (250 μM) for 30 min (o), 60 min (p) or 120 min (q) as indicated. **r** The viability of HeLa cells transfected with

TDP-43-HA together with vector or Hsp70s, treated with arsenite (100 μM) for indicated durations is assessed using the CCK-8 assay. Mean ± SEM; n = ~100 cells in (m-n) from pooled results of 3 independent repeats, n = 8 in (o-q), and n = 3 in (b, r). Student's *t* test (b, m and r) and two-way ANOVA (n-q); ns, not significant. Scale bars, 5 μm (c-l).

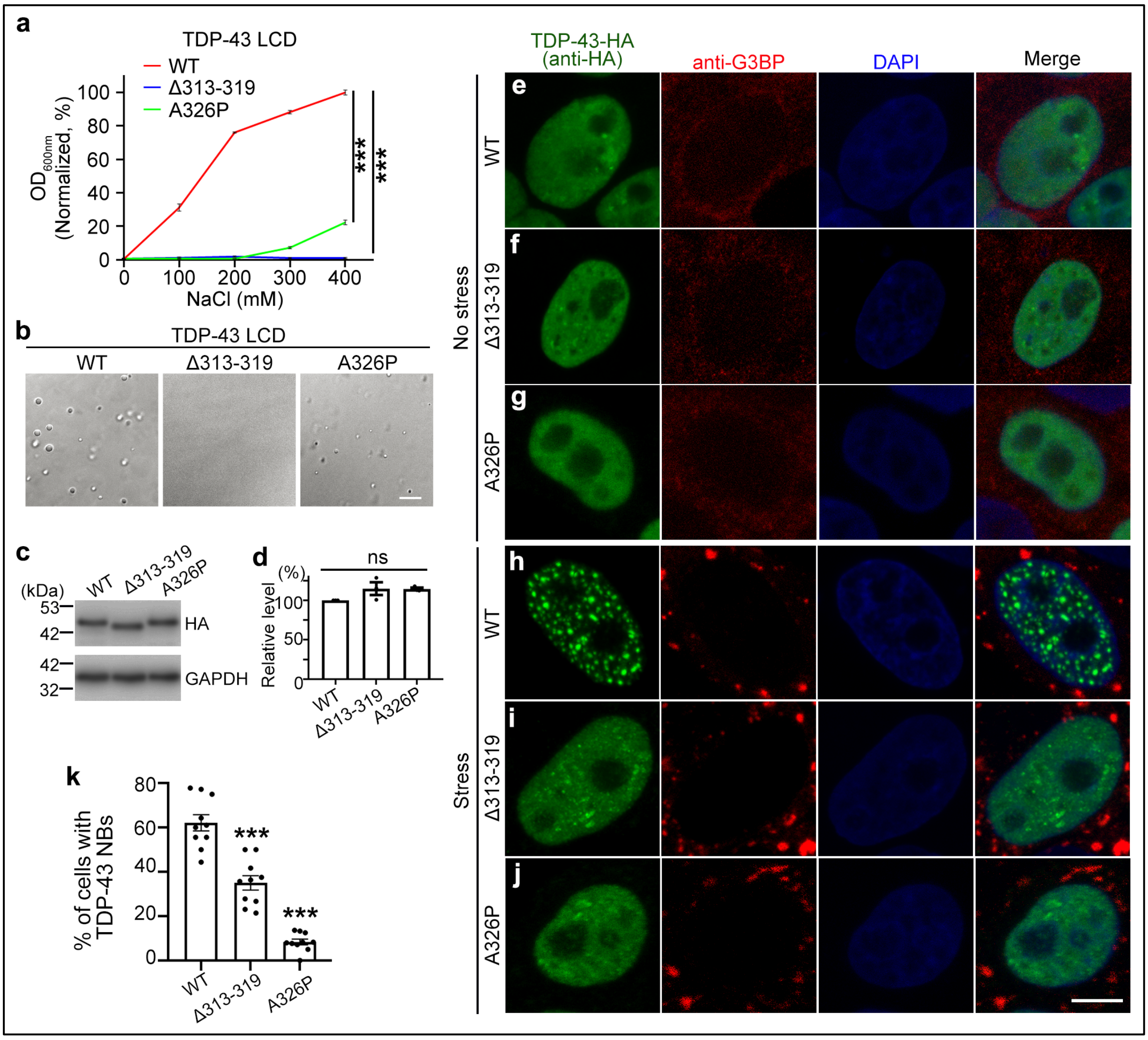


**Fig. S5: TDP-43-Δ313-319 and TDP-43-A326P exhibit severely impaired capability of forming liquid droplets *in vitro* and in cells.**

**a** Turbidity measurement of WT or the Δ313-319 and A326P variants of TDP-43 LCD (65 μM) with increasing concentrations of NaCl in 20 mM MES, pH 6.0. **b** DIC images of WT, Δ313-319 or A326P TDP-43 LCD (65 μM) in 400 mM NaCl, 20 mM MES, pH 6.0. **c-d** Representative images (c) and quantification (d) of Western blot analysis confirming the expression of WT or mutant TDP-43 FL in e-k. GAPDH is used as a loading control. **e-j** Representative images of HeLa cells expressing WT or mutant TDP-43 in the absence (e-g) or presence (h-j) of arsenite (250 μM, 30 min). DAPI, nuclear labeling; anti-G3BP for SGs. **k** The percentage of transfected cells showing TDP-43 NBs in h-j. Mean ± SEM, n = 3 in (a, d), and n = ~100 cells in (k) from pooled results of 3 independent repeats. Two-way ANOVA (a) and one-way ANOVA in (d, k); ***$p < 0.001$; ns, not significant. Scale bars, 10 μm (b) and 5 μm (e-j).
